# CSF1R inhibition during cranial radiotherapy reshapes the immune environment and glial dynamics

**DOI:** 10.1101/2025.10.11.681366

**Authors:** Yoran M Leter, Caroline L van Heijningen, Zoey J Tolboom, Mark C de Gooijer, Annette C Compter, Olaf van Tellingen, Sanne B Schagen, Laura E Kuil

## Abstract

**Background:** Cranial radiotherapy (cRT) induces long-lasting alterations in the brain microenvironment, including changes in glial populations and loss of neurogenesis. Pharmacological depletion of microglia via colony-stimulating factor 1 receptor (CSF1R) inhibition has been proposed as a strategy to modulate radiation-induced neuroinflammation, but its effects on other glial populations remains incompletely understood.

**Methods:** Nine-week-old male C57BL/6JRj mice (n=60) received either a CSF1R-inhibitor supplemented diet (pexidartinib, PLX3397) or a control diet prior to and continued for 10 days after fractionated CT-guided cRT (3x10 Gy, 30 Gy). Brain tissue was assessed at multiple time-points up to six months post cRT using immunohistochemistry for markers of microglia, astrocytes, oligodendrocytes and proliferating cells. Microglial identity and repopulation dynamics were assessed using IBA1, F4/80 and the homeostatic microglial marker P2RY12, complemented by morphometric analysis.

**Results:** Pexidartinib alone resulted in partial microglial depletion, whereas its combination with cRT led to near-complete loss of P2RY12+ homeostatic microglia at early time points. After discontinuation of the diet this was accompanied by increased IBA1+;F4/80^high^;P2RY12^neg^ cell densities, consistent with replacement of the microglial niche by monocyte-derived cells. Over time, these cells exhibited gradual morphological adaptation but remained largely P2RY12^neg^, indicating incomplete restoration of a homeostatic microglial phenotype. Combined treatment accelerated astrocytic reactivity compared to cRT alone and did not prevent cRT-induced loss of neurogenesis or OPCs. At later time points, partial OPC recovery was observed in selected regions in the combined treatment group.

**Conclusion:** CSF1R inhibition during cRT profoundly alters the composition and temporal dynamics of the brain’s myeloid compartment, characterized by sustained loss of homeostatic microglia and replacement by monocyte-derived cells. Under these experimental conditions, microglial depletion did not mitigate key histopathological features associated with radiation-induced brain injury and was associated with accelerated astrocytic responses. These findings highlight the importance of timing, extent of depletion, and repopulation dynamics when considering microglia-targeting strategies in the context of cranial radiotherapy.

**Key points:**

- Pexidartinib and cranial radiotherapy (cRT) have synergistic effects on microglial ablation
- Infiltrating monocytes repopulate the irradiated brain once the pexidartinib diet is discontinued
- In the absence of microglia, astrocytes show an accelerated reactivity to cRT
- Long-term after cRT, oligodendrocyte progenitor cell repopulation is enhanced in the pexidartinib treated animals

**Graphical abstract:** 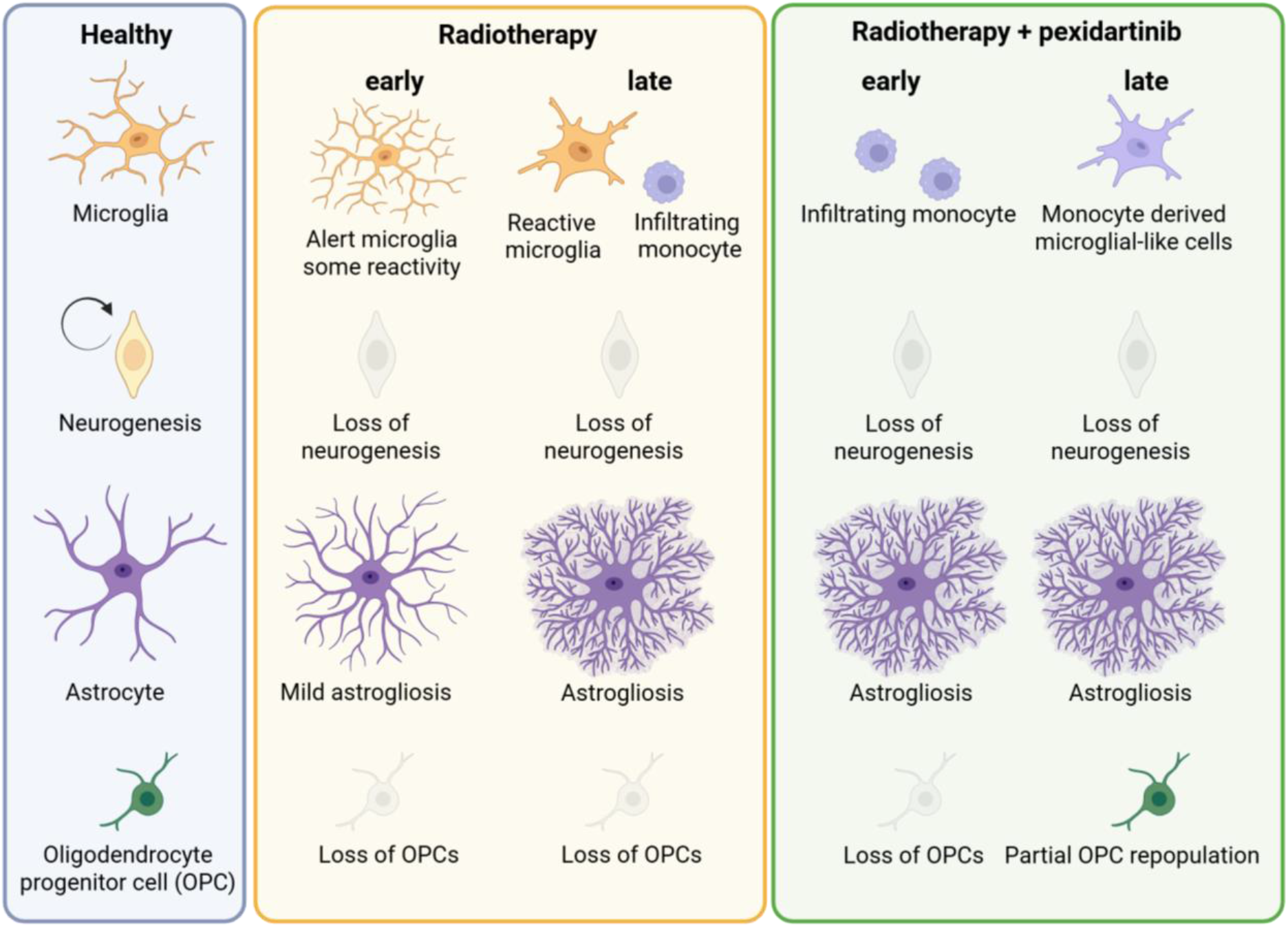

Generated by Biorender O.v.Tellingen (2026).

## Introduction

Cranial radiotherapy (cRT) is a common treatment for central nervous system (CNS) tumors such as primary brain tumors and brain metastases (Vogelbaum et al., 2022; Weller et al., 2023). Brain metastases (BM) occur in 10 to 40% of cancer patients, most frequently stemming from lung, breast, skin, kidney and gastrointestinal cancers (Rahman et al., 2022; Zhang et al., 2012). Up to 50% of CNS cancer patients treated with partial or whole-brain cRT develop progressive cognitive impairment (Cramer et al., 2019; Gonzalez-Johnson et al., 2024). This impairment, ranging from mild to profound and often progressive, substantially reduces patients’ quality of life by affecting cognitive performance and daily activities (Neo et al., 2017). Efforts to reduce cognitive problems upon cRT are undertaken including treatment with memantine and hippocampal sparing-RT, but in most cases this does not prevent cognitive impairment (Bou Dargham et al., 2025; Brown et al., 2020; Gondi et al., 2014). The cognitive impairment largely results from the neurobiological changes to the healthy brain tissue induced by radiation and there remains a critical need for interventions to prevent, reduce or revert.

Cognitive impairment following cRT are associated with demyelination of white matter, appearing as T2 hyper intensities on MRI in both pediatric and adult brain tumor patients (Connor et al., 2016; Ma et al., 2023; Marino et al., 2024; Voon et al., 2022). Preclinical studies implicate neuroinflammation in cRT-induced cognitive decline (Cramer et al., 2019). We and others showed neurobiological changes upon cRT including the loss of neurogenesis and oligodendrocyte progenitor cells (OPCs), astrocytic hypertrophy, reduced microglial numbers, and altered microglial morphology reflecting their reactive state (Han et al., 2016; Kuil et al., 2025; Monje et al., 2003; Panagiotakos et al., 2007; Voshart et al., 2024).

OPCs not only generate myelinating oligodendrocytes but also regulate microglia, neuronal circuitry, vascularization, and blood–brain barrier (BBB) integrity (Kimura et al., 2020; Pepper et al., 2018). Upon myelin damage, OPCs respond rapidly and play a role in the replenishment of myelination (Franklin & Ffrench-Constant, 2017). Thus, cRT–induced OPC loss may contribute to the late-delayed demyelination observed in treated patients (Wiggermann et al., 2019). However, ways to protect the OPCs from radiotherapy-induced death or stimulate their repopulation are absent.

Microglia spring from primitive macrophages produced during embryonic yolk sac primitive hematopoiesis and maintain their population by self-renewal throughout life (Askew et al., 2017; Masuda et al., 2022). After BBB maturation, microglia remain confined to the CNS, and peripheral macrophages are excluded, unless barrier integrity is compromised (Reddaway et al., 2023). Microglia express markers such as IBA1, P2RY12, TMEM119, F4/80, and CX3CR1, with loss of some homeostatic markers (P2RY12, CX3CR1, TMEM119) indicating reactive states (De Simone et al., 2010; Paolicelli et al., 2022). In their homeostatic state, microglia display a small soma with long, highly branched processes that continuously survey the environment. Upon detection of cell injury or death, they adopt a reactive morphology characterized by an enlarged soma and thick, shortened branches (Reddaway et al., 2023). In severe pathological conditions, including neurodegenerative diseases, infection, stroke, tumors, brain injury, and neuroinflammation, microglia can fully retract their branches, assuming an amoeboid form resembling peripheral macrophages in their morphological appearance (Nayak et al., 2014; Reddaway et al., 2023).

Temporary depletion of microglia via colony-stimulating factor 1 receptor (CSF1R) inhibition, followed by rapid repopulation, has been shown in preclinical neurological disease models to reduce neuroinflammation, restore microglial morphology, and prevent memory problems (Dagher et al., 2015; Henry et al., 2020; Iba et al., 2025; Thi Lai et al., 2024). In cRT models, persistent microglial activation is linked to cognitive dysfunction, and eliminating microglia during and after cRT using the CSF1R inhibitor PLX5622 prevented these deficits in mice (Acharya et al., 2016; Feng et al., 2021; Feng et al., 2016; Feng et al., 2018). These findings have led to the prevailing hypothesis that removing microglia during and after cRT improves brain health. However, a recent study showed no protection by CSF1R inhibitor treatment against cRT-induced loss of neurogenesis (Zhou et al., 2025), thus not all negative consequences of cRT might be prevented. Also, most studies have focused on short-term cognitive outcomes and microglia-specific endpoints, leaving it unclear how transient microglial ablation affects other vulnerable cell populations or the longer-term glial landscape after cRT.

Modulating microglia through CSF1R inhibition during cRT as a strategy to modulate microglia-dependent injury responses is conceptually attractive, but the integrated response of the irradiated CNS is likely multifaceted. In this study, we aimed to characterize the neurobiological long-term effects of CSF1R inhibition in the context of fractionated cranial radiotherapy using a dose regimen relevant to high-dose clinical scenarios. By initiating CSF1R inhibition prior to irradiation, we specifically sought to examine how a microglia-depleted or microglia-compromised environment influences the subsequent glial and myeloid response to radiation. We performed a longitudinal analysis of microglial markers, including the homeostatic marker P2RY12, alongside broader myeloid markers, assessed astrocytes, OPCs and neurogenesis across multiple brain regions in mice.

Our findings reveal that CSF1R inhibition in this setting leads to a sustained loss of homeostatic microglia and a marked shift in the composition of the brain’s myeloid compartment, accompanied by altered astrocytic responses and limited effects on radiation-induced depletion of neural and glial precursor populations. These results highlight the context-dependent nature of microglia-targeting strategies and underscore the importance of considering radiation dose, timing of intervention, and repopulation dynamics when evaluating their potential impact on the irradiated brain.

## Results

### Synergistic microglial loss with pexidartinib + cRT

To assess microglial sensitivity to cRT in the presence or absence of CSF1R inhibition, mice were fed a pexidartinib-supplemented or control diet during three weeks prior to cRT, continuing until 10 days after treatment (Fig 1A). A first important observation is that although 290mg Pexidartinib per kg diet was supplemented, this resulted in 60% lower plasma levels compared to the study by Elmore et al (Elmore et al., 2014)(Fig 1B). Sham-treated animals on pexidartinib diet displayed reduced IBA1+ cells in cortex, hippocampus, and thalamus (41.3% reduction in cortex, 25.8% reduction in hippocampus, and 85.6% reduction in thalamus, Fig 1C-D). Intriguingly, on top of this modest depletion, combining pexidartinib with cRT produced a near-complete loss of IBA1+ microglia across regions (95.4% cortex, 95.8% hippocampus, 85.6% thalamus; all p<0.05), suggesting enhanced radiosensitivity of pexidartinib-treated microglia as pexidartinib levels in the brain were similar in cRT + pexidartinib and pexidartinib only animals (Fig 1C-D). Thus, CSF1R inhibition partially depleted microglia, but sensitized them to cRT. Therefore, microglia were synergistically ablated, resulting in the microglial ablation we aimed for in this experimental group.

**Figure 1.**
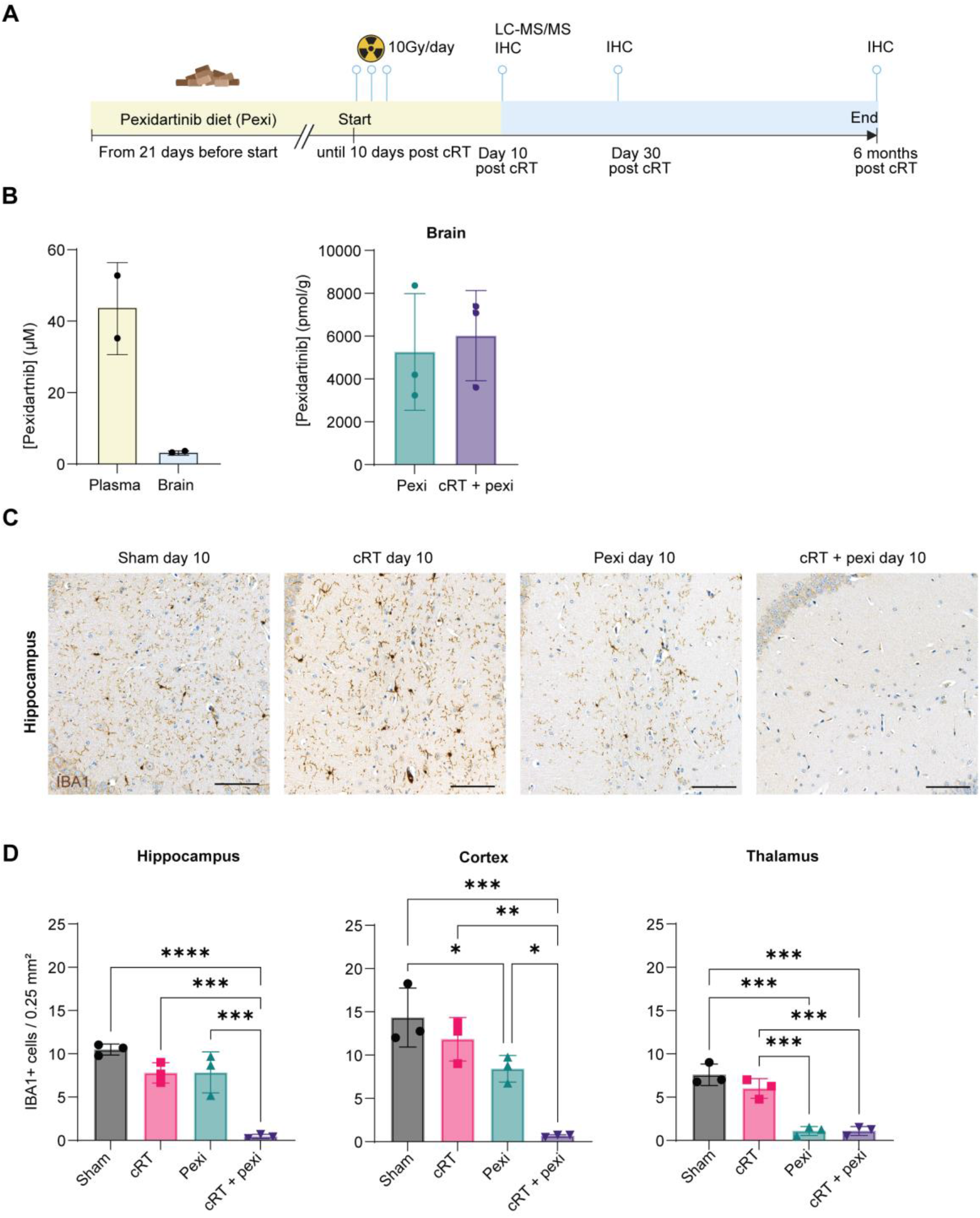
Experimental set-up of microglial ablation during cRT. A. Schematic representation of the experimental timeline where in yellow the pexidartinib diet (290mg/kg in chow) was supplemented starting 21 days preceding cRT continuously through 10 days post cRT. B. D. LC-MS/MS measurement of Pexidartnib levels in plasma and brain. C. IBA1 IHC in the hippocampus. D. Quantification of IBA1+ cells. Each dot represents one animal, error bars represent standard deviation. * p<0.05, ** p<0.01, ***p<0.001, ****P<0.0001.

### Monocyte-derived macrophage infiltration after microglial ablation

At 30 days post cRT treatment (20 days off CSF1R inhibitor diet), IBA1+ cells in the thalamus of cRT + pexidartinib treated animals were almost completely recovered, whereas IBA1+ cells in hippocampus and cortex of cRT and pexidartinib standalone treatment groups showed no change relative to day 10 (Fig 1C *vs.* 2A and S1-2). cRT + pexidartinib animals showed significantly higher IBA1+ cell numbers, with a clearly different morphology, than cRT alone in cortex, hippocampus, and thalamus (p<0.05)(Fig 2A; S1-2). IBA1 is a PAN-macrophage marker known to label microglia, but also CNS-resident macrophages and peripheral macrophages. Therefore, P2RY12 was used as a marker expressed by homeostatic microglia, but not expressed by peripheral or CNS-associated macrophages. It is important to note that P2RY12 is downregulated upon microglial reactivity (Paolicelli et al., 2022). P2RY12 staining revealed an almost complete loss of homeostatic microglia in the cRT + pexidartinib group, while cRT alone showed just a partial loss of homeostatic microglia (Fig 2A; S1-2). We, next, quantified F4/80+ cells and distinguished between F4/80^high^ monocytes and F4/80^low^ microglia. This revealed widespread monocyte engraftment in cRT + pexidartinib animals, evident from the distinct population of intensely stained F4/80^high^ cells compared to the other groups (Fig 2A; S1-2). This difference was quantified using Qupath’s automated H-scoring based on three staining intensity thresholds, resulting in a significantly higher H-score in the cRT + pexidartinib group (Fig S1-2). This result is consistent with previous work showing that the microglial niche, emptied by CSF1R inhibition, is competitively repopulated by proliferating F4/80^low^Clec12a^-^ microglia and F4/80^high^Clec12a^+^ infiltrating monocytes (Bastos et al., 2025; Lund et al., 2018).

**Figure 2.**
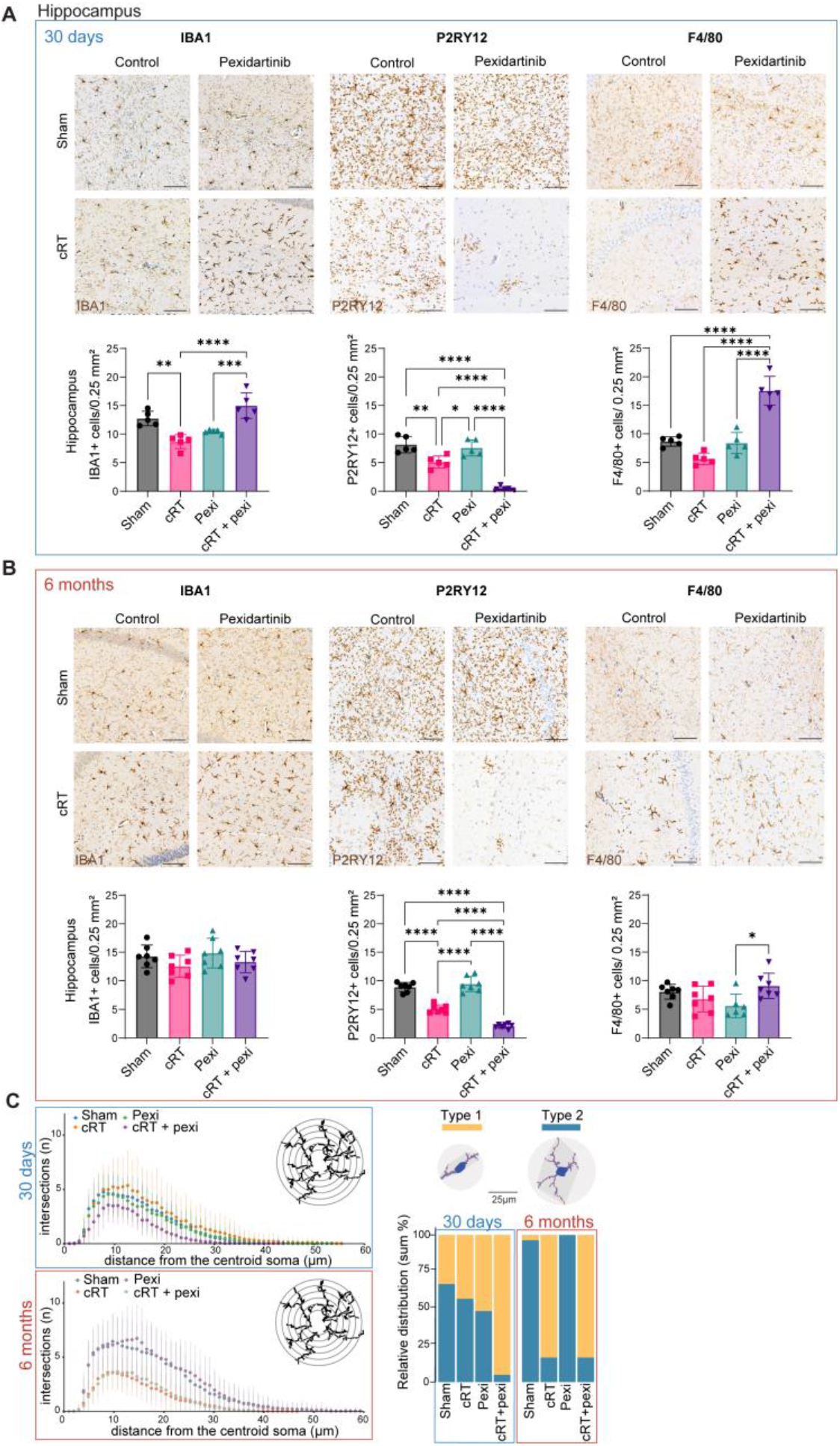
Quantification and morphological assessment IBA1 cells. A. IHC of IBA1, P2RY12 and F4/80 in the hippocampus at 30 days post cRT. B. IHC of IBA1, P2RY12 and F4/80 in the hippocampus at 6 months post cRT. Each dot represents one animal, error bars represent standard deviation. * p<0.05, ** p<0.01, ***p<0.001, ****P<0.0001. C. Morphometric analysis using mGlia showing a sholl plot and relative cluster distribution plot.

### Reduced microglial numbers persist half a year post cRT in cortex and thalamus

At 6 months post cRT, IBA1, P2RY12 and F4/80 positive cell numbers were still reduced in the cortex and thalamus in the cRT only group (Fig S1-2). This confirms our earlier observation that cRT reduces microglial densities (Kuil et al., 2025). In the hippocampus the number of P2RY12+ homeostatic microglia remained decreased after cRT only, whereas IBA1 and F4/80 cell numbers returned to baseline levels (Fig 2B). In the cRT + pexidartinib animals the number of P2RY12+ homeostatic microglia was still significantly decreased although there is some recovery relative to day 30, especially in cortex (Fig 2B; S1-2).

### Morphology of IBA1 cells upon cRT is similar regardless of pexidartinib treatment at 6 months

Morphometric analyses were performed on cells in the hippocampus to characterize the morphological differences induced by cRT and/or pexidartinib at 30 days and 6 months post cRT. At 30 days, the sham and pexidartinib groups contained respectively 33% and 52% of cells presenting a type 1 morphology (less branched cells). At 6 months, both groups showed almost exclusively cells with a type 2 morphology (high branching complexity)(Fig 2C). Thus, upon aging the complexity of IBA1+ microglia in the brain increases and this is not changed by pexidartinib treatment. In the cRT-only group cells shifted from a mixed type 1/2 morphology towards a type 1 morphology between 30 days and 6 months post cRT. The cRT + pexidartinib treated animals were already predominantly type 1 at day 30 post treatment. By 6 months, there was a slight proportional conversion back to a type 2 morphology, resulting in a net distribution between type 1 and type 2 morphologies similar to the cRT only group (Fig 2C). This decreased branching complexity was observed regardless of pexidartinib treatment (p<0.0001 for both groups)(Fig 2C). Intriguingly, this shows that the morphology of IBA1 cells in the two groups become more similar over time. This suggests that in the cRT only condition, the IBA1+ cells in the brain might, over the following months, include not only reactive microglia (P2RY12^neg^) but also some infiltrating monocytes (F4/80^high^P2RY12^neg^). At the same time, cRT + pexidartinib IBA1+ cells represent infiltrating monocytes that adapt a microglia-like morphology over time, increasing the relative proportion of cells with a type 2 morphology at 6 months post cRT compared to 30 days post cRT.

### cRT-induced effects on oligodendrocyte progenitor cells (OPCs), myelination, neurogenesis and astrogliosis

OPCs (PDGFRA+) were significantly reduced in hippocampus and cortex at 10 days after cRT, which was unaffected by pexidartinib co-treatment (Fig 3A; S3A). This depletion persisted at 30 days, when also the thalamus showed OPC ablation (Fig 3A; S3A). Interestingly, partial OPC recovery occurred in cRT + pexidartinib animals in hippocampus and thalamus at 6 months post cRT, while in the cRT alone animals OPCs no recovery over time was observed (Fig 3A; S3A).

**Figure 3.**
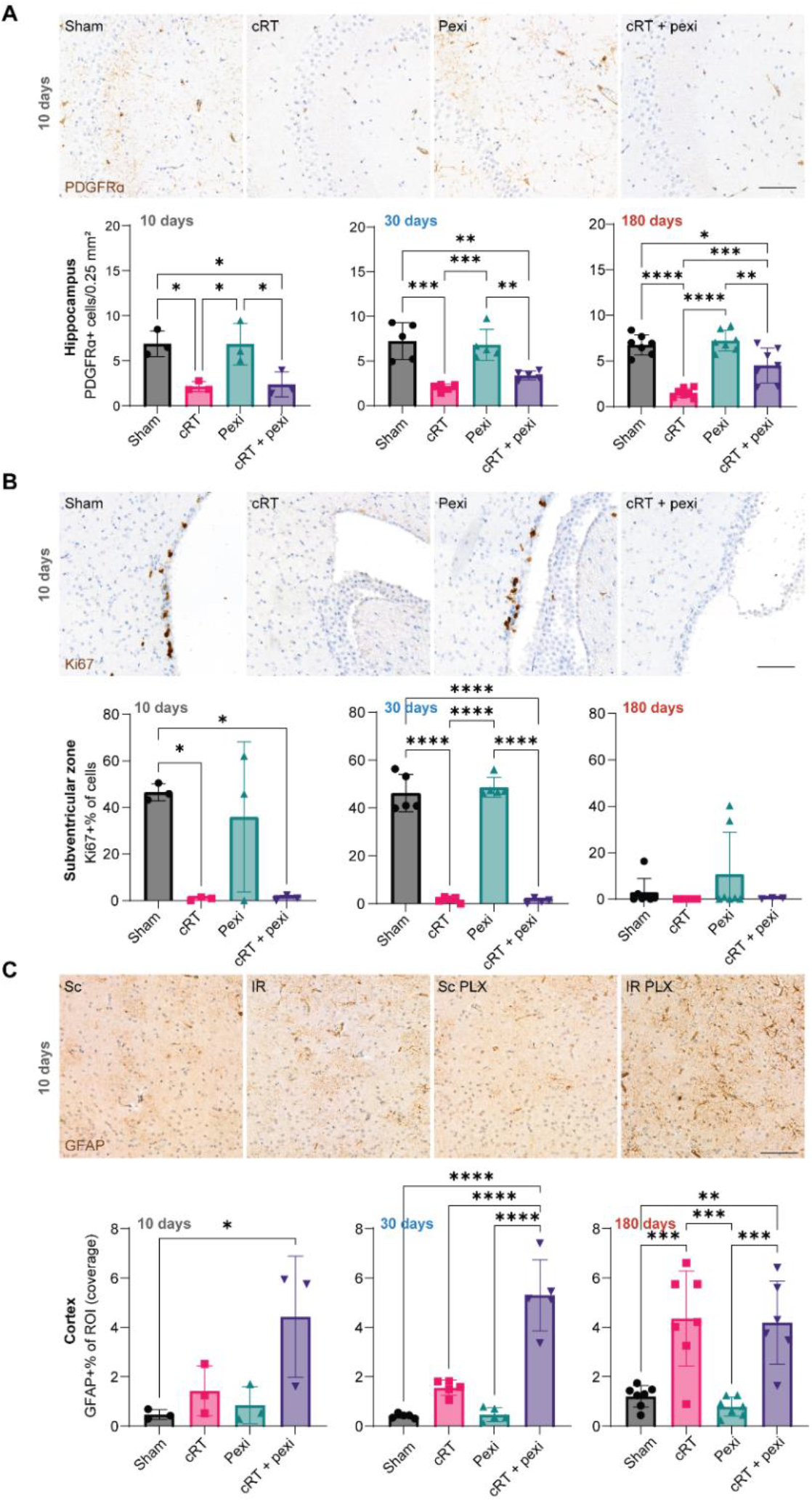
IHC analysis of OPCs, neurogenesis and astrocytes. A. IHC of PDGFRA labelling OPCs in the hippocampus at 10 days post cRT and quantification of PDGFRA+ cells on all three time points B. IHC of Ki67 labelling proliferation in the sub-ventricular zone (neurogenic site) at 10 days post cRT and quantification of Ki67+ cells normalized to total cells on all three time points. C. IHC of GFAP labelling astrocytes in the cortex at 10 days post cRT and quantification of GFAP+ pixel coverage normalized to total region area on all three time points. Each dot represents one animal, error bars represent standard deviation. * p<0.05, ** p<0.01, ***p<0.001, ****P<0.0001.

Myelin basic protein (MBP) coverage, reflecting myelination, was mildly reduced by cRT in the parenchyma of the cortex (n.s.) with no rescue by pexidartinib (Fig S4A). The MBP coverage of the perforant pathway, the white matter tract connecting the entorhinal cortex to the hippocampal formation, was significantly reduced upon cRT and was not rescued by pexidartinib treatment (Fig S4A).

Ki67+ proliferative cells in hippocampal neurogenic regions, reflecting neural stem cell proliferation, were nearly absent after cRT at each time point, regardless of pexidartinib treatment (Fig 3B; S3B). Thus, pexidartinib treatment did not rescue the cRT-induced loss of neurogenesis.

GFAP+ coverage increased after cRT in multiple regions, reflecting reactive astrocytes (Fig 3C; S4B). At early time points (10–30 days), this response was significantly amplified by cRT + pexidartinib in cortex and medial prefrontal cortex, but not in the hippocampus (Fig 3C; S4B). By 6 months, GFAP coverage was similarly elevated in both irradiated groups, reflecting accelerated gliosis in the cRT + pexidartinib group compared to cRT alone (Fig 3C; S4B).

## Discussion

Cranial radiotherapy (cRT) triggers complex, long-lasting changes in the brains glial landscape. We show that transient CSF1R inhibition with pexidartinib during cRT synergistically depletes microglia, drives rapid and widespread engraftment of monocyte-derived macrophages, and accelerates astrocytic reactivity, yet fails to protect neural stem cells or oligodendrocyte progenitor cells (OPCs). Over time, infiltrating monocytes adopt microglia-like morphologies and seems to favor OPC repopulation. However, neither myelin integrity nor neurogenesis recovers, underscoring the limited neuroprotective potential of this strategy.

### Synergistic microglial depletion upon cRT + pexidartinib

Our study shows that mild microglia depletion induced by CSF1R inhibition is aggravated by cRT in a synergistic manner. Previous studies have shown that another CSF1R inhibitor, GW2580, does not reduce microglial numbers but instead sensitizes microglia to reactive oxygen species, which might make them more sensitive to cRT (Soto-Diaz et al., 2021). Another possible explanation for the increased microglial radiosensitivity is that pexidartinib-induced microglia loss triggers proliferation in the remaining microglia, thereby increasing the number of microglia in mitosis during cRT. It is well known that cells in mitosis are more sensitive to DNA-damaging cRT (Cosper et al., 2023; Stobbe et al., 2002).

### Empty microglial niche allows for infiltration of the brain by monocyte-derived cells

Our findings challenge the prevailing assumption that microglial ablation during cRT is broadly beneficial. Previous rodent studies reported improved cognition after CSF1R inhibition combined with lower-dose cRT (Acharya et al., 2016; Feng et al., 2018), but these studies primarily assessed short-term outcomes and focused on microglial reactivity and dendritic complexity without assessing effects on other glia. By extending the observation window to six months and examining multiple glial lineages, we demonstrate that while pexidartinib intensifies early microglial loss, it also creates an empty niche permissive to monocyte infiltration. Consistent with lineage-tracing studies using lower-dose cRT and CSF1R inhibitors (Bastos et al., 2025; Du et al., 2024; Feng et al., 2021; Hohsfield et al., 2020; Morganti et al., 2014; Xu et al., 2020), these infiltrating cells persist long-term, suggesting stable engraftment.

Zhou and colleagues (2025) showed that depletion of microglia delayed the removal of dead neuronal progenitors and strongly increased CCL2 levels, resulting in a pro-inflammatory environment (Zhou et al., 2025). A pro-inflammatory environment could have persistent negative effects, as pro-inflammatory states have been found to negatively influence microglia migration towards the ADP gradient, a process modulated by the P2RY12 receptor and essential for microglial immune response (De Simone et al., 2010). Elevated CCL2 levels likely induced infiltration of monocytes under these conditions, given that monocytes can be attracted to the brain by CCL2-CCR2 signaling (Howe et al., 2017; Mondini et al., 2019; Murugan et al., 2020; Varvel et al., 2016). This seems replicated in our model, as shown by the infiltration of F4/80^high^ cells and the absence of P2RY12+ cells in the cRT + pexidartinib animals 30 days post cRT and the abundance of reactive astrocytes 10 days after cRT in the cRT + pexidartinib animals that persists over time. Feng et al. used a similar set-up to ours and showed that replacement of microglia by monocytes can be protective at the level of synapses and cognition (Feng et al., 2021). Our work shows that, at the glial lineage level, this replacement is not broadly protective and has complex, cell-type–specific consequences.

### Brain-engrafted monocyte-derived cells become microglia-like over time

Assessing the morphology of IBA1+ cells at 30 days after treatment showed increased branching complexity following cRT, as well as in the sham-treated animals over time. These findings are in line with our previous observations in early-aging (77 weeks) female animals and with previously reported morphological changes after cRT in male animals (Kuil et al., 2025). Based on the changes in IBA1+ cell morphology in the cRT + pexidartinib group over time (between 30 days and 6 months) and the mild increase in P2RY12+ homeostatic microglial cell numbers, one might speculate that the infiltrating monocyte-derived microglia gradually acquire a microglial-like identity. This has been suggested by Feng and colleagues following cRT, as well as in other contexts, such as stroke, lesions, and models for multiple sclerosis (Chen et al., 2020; Feng et al., 2021; Mildner et al., 2007). However, the question of whether these cells truly can become microglia in terms of transcriptome and identity remains unresolved (Bastos et al., 2025; Cronk et al., 2018). Furthermore, it is still unclear whether monocyte-derived cells are beneficial, detrimental, or neutral for long-term brain health under normal conditions or after cRT.

### Microglial ablation during and after cRT did not prevent NSC and OPC loss

Microglial ablation during cRT did not prevent the loss of neurogenesis, an early and persistent effect of cRT. In line with our observations, Zhou et al. showed that cRT-induced loss of neurogenesis occurs regardless of microglial presence, using a genetic microglial ablation model in juvenile mice (Zhou et al., 2025). This challenges the general perception that microglia are detrimental to neurogenesis following cRT due to their reactivity (Gibson & Monje, 2021; Monje et al., 2003; Voshart et al., 2024).

cRT-induced loss of OPCs was also not prevented by pexidartinib treatment. This contrasts with findings from Gibson et al., who reported that CSF1R inhibition prevented chemotherapy-induced OPC loss (Gibson et al., 2019). Interestingly, long-term post cRT, OPCs showed enhanced repopulation in pexidartinib + cRT animals compared to cRT only animals. This suggests that the microenvironment induced by pexidartinib treatment in combination with cRT is, over time, beneficial for OPC repopulation. This effect may result from the presence of monocyte-derived macrophages in the brain instead of endogenous (reactive) microglia, or from an accelerated immune response consisting of reactive macrophages and reactive astrocytes that is initiated already early after cRT.

### CSF1R inhibitor treatment during cRT induced an accelerated astrocyte reactivity

A seminal *in vitro* study by Liddelow et al. showed that activated microglia release IL-1α, TNF, and C1q, which are necessary to induce the formation of neurotoxic astrocytes (Liddelow et al., 2017). Along the same lines, Gibson et al. showed that chemotherapy on itself did not induce reactivity of astrocytes *in vitro* using monocultures (2019). However, in the presence of medium of chemotherapy-induced reactive microglia, reactive astrocytes were induced (Gibson et al., 2019). Unfortunately, they did not assess astrocytes in their *in vivo* CSF1R inhibitor + chemotherapy model. In contrast, our data show that microglial presence is not required for astrocyte reactivity upon cRT, as CSF1R-induced ablation of microglia accelerated the astrocytic reactivity to cRT. Rapid responses of both microglia and astrocytes upon lipopolysaccharide (LPS) model triggering immune reactions or experimental autoimmune encephalitis (EAE) model triggering myelin damage have been observed in previous *in vivo* studies (Chanaday & Roth, 2016; Xingi et al., 2023). However, studies that tried to induce reactive astrogliosis in the absence of microglia are limited. One study showed that minocycline did not prevent astrocytic reactivity in response to stroke (Bourget et al., 2022). However, in their additional experiment using CSF1R inhibitor treatment, stroke itself did induce astrocytes, but the effect did not reach significance, hampering the assessment of a possible improvement with CSF1R inhibition (Bourget et al., 2022). Thus, the complex interactions between microglia and astrocytes remain elusive, but in the absence of microglia, our cRT model induces a major astrocytic response that is sustained over time.

### Timing and extend of microglial ablation and repopulation might be critical

In future research, it might be valuable to alter the timing of microglia depletion relative to the administration of cRT. For example, Henry et al. (2020) depleted microglia one month after traumatic brain injury, followed by quick repopulation, allowing microglia to resume their homeostatic function shortly after injury (Henry et al., 2020). This approach could be used to investigate whether pro-inflammatory conditions following cRT can be prevented by microglia depletion and how this affects long-term cognitive outcomes. However, the study from Hohsfield et al. (2020) suggests that even after BBB recovery for 3.5 months after cRT, CSF1R inhibitor-induced microglial ablation still results in infiltration of peripheral monocytes instead of endogenous microglial expansion (Hohsfield et al., 2020). To prevent infiltration of monocytes, one might adjust the CSF1R dosing to induce only partial ablation, resulting in higher remaining microglial numbers in the brain that might prevent monocyte infiltration and allows endogenous microglia to repopulate the partially empty niche. This partial ablation strategy has been shown to normalize microglial primed state towards resting state (Leon-Rodriguez et al., 2025) and in aged mice to improve synaptic connectivity and cognitive function (Strackeljan et al., 2025).

### Conclusion

This study demonstrates that CSF1R inhibition during cranial radiotherapy substantially reshapes the brain’s glial landscape, leading to a prolonged reduction of P2RY12+ homeostatic microglia and repopulation by IBA1+ myeloid cells lacking canonical microglial markers. These findings suggest that microglial depletion in this setting does not result in restoration of a homeostatic microglial compartment, but rather in a persistent shift in myeloid cell identity.

In parallel, combined CSF1R inhibition and radiotherapy did not prevent loss of neurogenesis or oligodendrocyte progenitor cells and was associated with an accelerated astrocytic response at early time points. Although partial recovery of OPCs was observed at later stages in selected regions, broader markers of tissue integrity were not improved.

Together, these data indicate that microglial depletion during cranial radiotherapy leads to complex and context-dependent changes in the neuroimmune environment, underscoring the need for careful consideration of treatment timing and microglial repopulation dynamics in future therapeutic strategies.

## Material and methods

### Animals

Experiments were approved by the Animal Experimentation Committee of the Netherlands Cancer Institute (NKI) and all were carried out in accordance with the approved protocols. The principles of laboratory animal care, Dutch law and the guidelines for care and the Code of Practice of lab animals used in oncology were followed. The number of mice per group were determined based on previous experience with immunohistochemistry analysis. All methods are reported in accordance with ARRIVE guidelines. Five-week-old male (C57BL/6JRj) mice ordered from an external company (Janvier-Labs) were housed with 3-5 animals in clear cages (n=60). The cages contain wood shavings, food and water. Mice were exposed to a fixed 12:12 hour light:dark cycle. A visual representation of the experimental setup is shown in Figure 1B. At day -21 the experiment started with 10-week-old animals. Half of the mice received a regular diet (control)(n = 30) and half the pexidartinib (PLX3397) supplemented diet (290 mg/kg resulting in approximately 40 mg/kg/animal/day of intake, SAFE laboratories, France) (n = 30). 21 days post-diet start (day 0) mice were divided in 4 groups of 15 animals; sham control, irradiated (cRT), sham control pexidartinib diet (pexi) and irradiated pexidartnib diet (cRT + pexi). Mice were sedated with isoflurane (2-4%). Ct-guided cRT was administered with a X-RAD SmART (Small Animal Irradiator) using a s10 collimator. Mouse brains were irradiated with 5Gy bilaterally (90° and 270°)(total 10Gy/day) for 3 consecutive days (total dose of 30Gy). 30Gy irradiation is equal to an Equivalent Dose in 2 Gy Fractions (EQD2) of 90Gy (α/ß ratio: 2). After cRT, the mice were left to recover on a heated platform. The sham-irradiated animals were also subjected to this procedure (anesthesia), but without being exposed to Ct or irradiation. The diets continued until day 10 after cRT, when the first cohort of animals (n = 12, n = 3/group) was sacrificed for immunohistochemistry (IHC) analysis of PLX3397-induced microglia ablation and LC-MS/MS analysis of plasma and brain pexidartinib levels. From day 10 onwards all animals were put on regular control diet. Animals were sacrificed at 30 days post cRT (n = 20, n=5/group) and 6 months post cRT (n = 28, n=7/group) for IHC analysis of microglia presence and morphology (Figure 1). Animals were sacrificed by CO2 asphyxiation.

### LC-MS/MS

Liquid chromatography with tandem mass spectrometry (LC-MS/MS) was used to determine the plasma and brain levels of pexidartinib (MRM: 418.2 → 258,0) using buparlisib (411.0 → 367.1) as internal standard, The LC-MS/MS system consisted of an UltiMate 3000 LC System (Thermo Scientific, Waltham, MA, USA) and Triple Quad™ 3500 (SCIEX, MA, USA)). Data acquisition and quantification were carried out with Analyst 1.7.2 (Sciex) and LC instrument control using Chromeleon 7.2 (Thermo).

### Sample pretreatment

Calibration standard ranging from 5 nM to 10,000 nM pexidartinib, blank human plasma (Sanquin, Amsterdam, NL) and blank brain tissue homogenates. Six volumes of internal standard (IS) buparlisib (100 nM in Acetonitrile:Formic acid (99:1 v/v)) was added to one volume of sample while vortex-mixing. Next, samples were centrifuged (5 min, 20.000 g, 4 °C) and the supernatant was diluted in water (1:5 v/v) in a 96 wells plate (Thermo Fisher Scientific, CA, USA). Blanks (IS with blank matrices (blank human plasma or blank tissue) and double blanks (only blank matrices) were included in each measurement run. Samples of 50 µL were injected on a Symmetry C18 Column (3.5 µm, 2.1 mm X 150 mm; Waters Corporation, MA, USA) maintained at 50°C. Elution was achieved using a 1.5 minute gradient from 10% to 95% B (mobile phase (A): 0.1% formic acid in water (v/v), mobile phase (B): methanol) with a 0.4 mL/min flow. The 95% B was maintained for 3.5 minutes followed by re-equilibration at 10% B at a 0.6 mL/min flow for 4 min.

### Immunohistochemistry staining

Mouse brains were fixed in 4% formaldehyde for 48 hours at room temperature and embedded in paraffin by the animal pathology core facility of the Netherlands Cancer Institute. All slides were stained with DAB chromogen antibodies, with slight differences for each staining, highlighted in table 1. Secondary antibody EnVision+ System-HRP labelled Polymer a-rabbit, Dako K4003. Slides are scanned on a Panoramic P1000 slidescanner.

**Table 1.** Antibody details used for immunohistochemistry.

| Marker | Antigen retrieval | Pre-incubation | Incubation with primary antibody |
| --- | --- | --- | --- |
| **IBA1** | Citrate buffer (SCytek; CBB999) in pressure cooker at 110°C for 20 minutes followed by 30-minute cooldown time | 10% milk powder in PBS | IBA1 (FujiFilm; 019-19741) 1:2000 in PBS/1% BSA/1.25% NGS |
| **F4/80** | Tris/EDTH pH 9.0 (Tris (Sigma; 252895), EDTH (Sigma; EDS)) in PT module at 98°C for 20 minutes | PBS/4% BSA/5% NGS | F4/80 (Cell signaling; #70076s) 1:1000 in PBS/1% BSA/1.25% NGS |
| **P2Y12** | Tris/EDTH pH 9.0 (Tris (Sigma; 252895), EDTH (Sigma; EDS)) in pressure cooker at 110°C for 20 minutes followed by 30-minute cooldown time | PBS/4% BSA/5% NGS | P2Y12 (Anaspec; AS-55043A) 1:100 in PBS/1% BSA/1.25% NGS |
| **Ki67** | Tris/EDTA pH9.0 (Tris (Sigma; 252895), EDTH (Sigma; EDS)) in a PT module at 98°C for 20 minutes | PBS/4% BSA/5% NGS | Ki67 (anaspec as-55043A) 1:100 PBS/1% BSA/1.25% NGS |
| **PDGFRA** | Tris/EDTA pH9.0 (Tris (Sigma; 252895), EDTH (Sigma; EDS)) in a PT module at 98°C for 20 minutes | 10% milk powder in PBS | PDGFRA (Cell signalling 3169) 1:50 PBS/1% BSA/1.25% NGS |
| **MBP** | Citrate buffer (SCytek; CBB999) in pressure cooker at 110°C for 20 minutes | PBS/5% NGS | MBP (BioLegend; 836504) 1:500 in PBS/1% BSA/1,25% NGS |
| **GFAP** | Tris/EDTA (pH9.0) in water bath 96°C for 30 minutes, cooldown 30 minutes | 10% milk powder in PBS | GFAP (Biotrend; BT46-5002-04) 1:1000 in PBS/1% BSA/1,25% NGS |

### Manual cell count

For the manual cell count in mouse cortex and thalamus, four 500 * 500 µm areas were randomly selected per brain area per animal using Qupath (V0.5.1). Images were exported to ImageJ counted and the average cell density per 0.25 mm^2^ was calculated. Hippocampal cell count was done by selecting the entire present hippocampus in the IHC image. Cells from the entire hippocampus were counted and the cell density per 0.25 mm^2^ was calculated based on the size of the ROI.

### Microglia morphology analysis

Microglia morphology was analyzed using a semi-automated microglial morphology pipeline (MGlia)(van Weering et al., 2023). This analysis was performed on the IHC images of the sacrificed cohorts at 30 and 180 days by randomly selecting 12 microglia per animal from the hippocampus (n = 20 animals at 30 days, n = 28 animals at 180 days). Cells were selected in QuPath, creating 500 * 500-pixel single-cell images. Images were loaded into MGlia where we manually adjusted thresholds to correctly select all cells distinguishing cell soma and branches. Resulting in Sholl analysis and morphometric features on which a hierarchical clustering on principal components (HCPC) was performed using R (v.4.4.1) according to the protocol described by van Weering (2023). AUC calculations were performed on the sholl data using the Linear Trapezoidal Method in R (4.4.1).

### Immunohistochemistry quantification of F4/80 expression using QuPath H-scoring

F4/80 immunohistochemistry was quantified using QuPath (version 0.5.1) on brightfield whole-slide images in the same 500 * 500 µm areas used for the manual quantification. Colour deconvolution was performed using an experimentally derived haematoxylin–DAB stain vector to separate the haematoxylin nuclear counterstain from the DAB chromogen signal corresponding to F4/80 immunoreactivity. Cell detection was performed using the positive cell detection algorithm with the following adjusted parameters: Detection image = Optical Density Sum; Background radius = 4 µm; Gaussian smoothing sigma = 1 µm; Minimum cell area = 5 µm²; Detection threshold = 0.15; Cell expansion = 3 µm. F4/80 positivity was determined using the mean DAB optical density measured within the nuclear compartment (Nucleus: DAB OD mean). Cells were classified into four staining categories using a three-threshold approach: Negative DAB OD < 0.2; F4/80low (1+): DAB OD 0.2–0.5; F4/80intermediate (2+): DAB OD 0.5–1.0; F4/80high (3+): DAB OD > 1.0. An H-score was calculated according to the standard formula: H-score=(1×%1+)+(2×%2+)+(3×%3+) resulting in a score ranging from 0 to 300.

### Statistical analysis

Graphpad Prism 9.3.0 was used for data visualization and analysis of manual cellcount data (Boston, MA). One-way ANOVA was used with Tukey’s multiple comparisons test combined with a Brown Forsythe test for equality of variance. In case of unequal variances, a Brown-Forsythe Welch ANOVA was performed. If the data were not normally distributed, a Kruskal-Wallis test was performed with a Dunn’s multiple comparisons test. Statistical significance was assumed when p < 0.05.

## Declarations

### Ethics approval

Experiments were approved by the Animal Experimentation Committee of the Netherlands Cancer Institute (NKI) and all were carried out in accordance with the approved protocols. The principles of laboratory animal care, Dutch law and the guidelines for care and the Code of Practice of lab animals used in oncology were followed.

### Consent for publication

All authors approved this manuscript for publication.

### Availability of data and materials

Data will be made available upon reasonable request by the corresponding author.

### Authors’ contributions

Conceptualization: SS, OvT, LK, AC. Formal analysis: ZT, LK, YL. Investigation: LK, MdG, CvH, YL, ZT. Writing – original draft: YL, LK, Writing – review & editing: all authors, Supervision: LK, OvT, SS.

## Acknowledgements

We would like to thank the animal pathology core facility and animal facility of the NKI for assistance.

