## Supplementary material for "CSF1R inhibition during cranial radiotherapy reshapes the immune environment and glial dynamics": Fig S

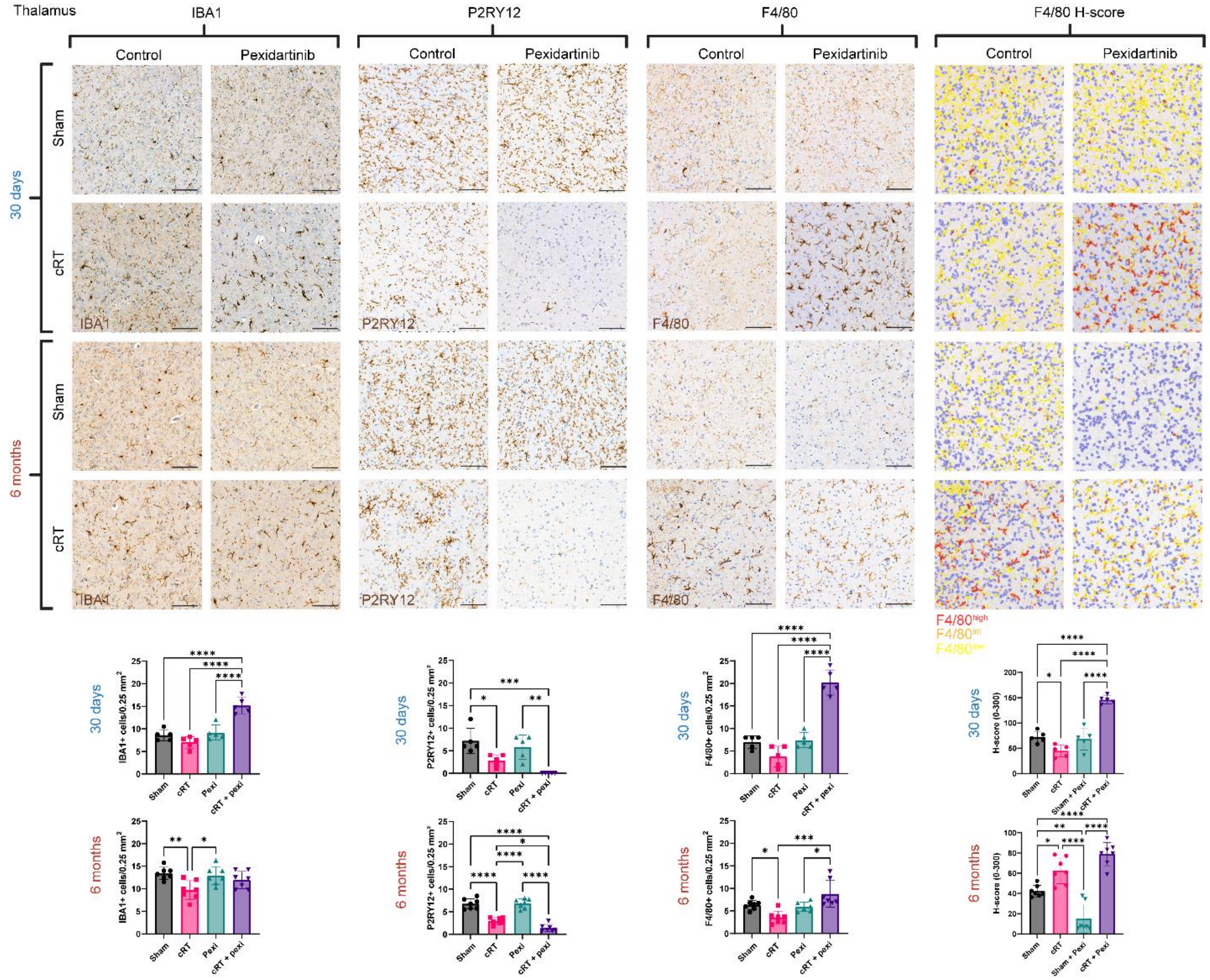


***Figure S1. IHC analysis of macrophage and microglial markers in the thalamus.*** *Representative images of IHC and quantification on all three time points in the thalamus. Each dot represents one animal, error bars represent standard deviation. * p<0.05, ** p<0.01, ***p<0.001, ****P<0.0001.*

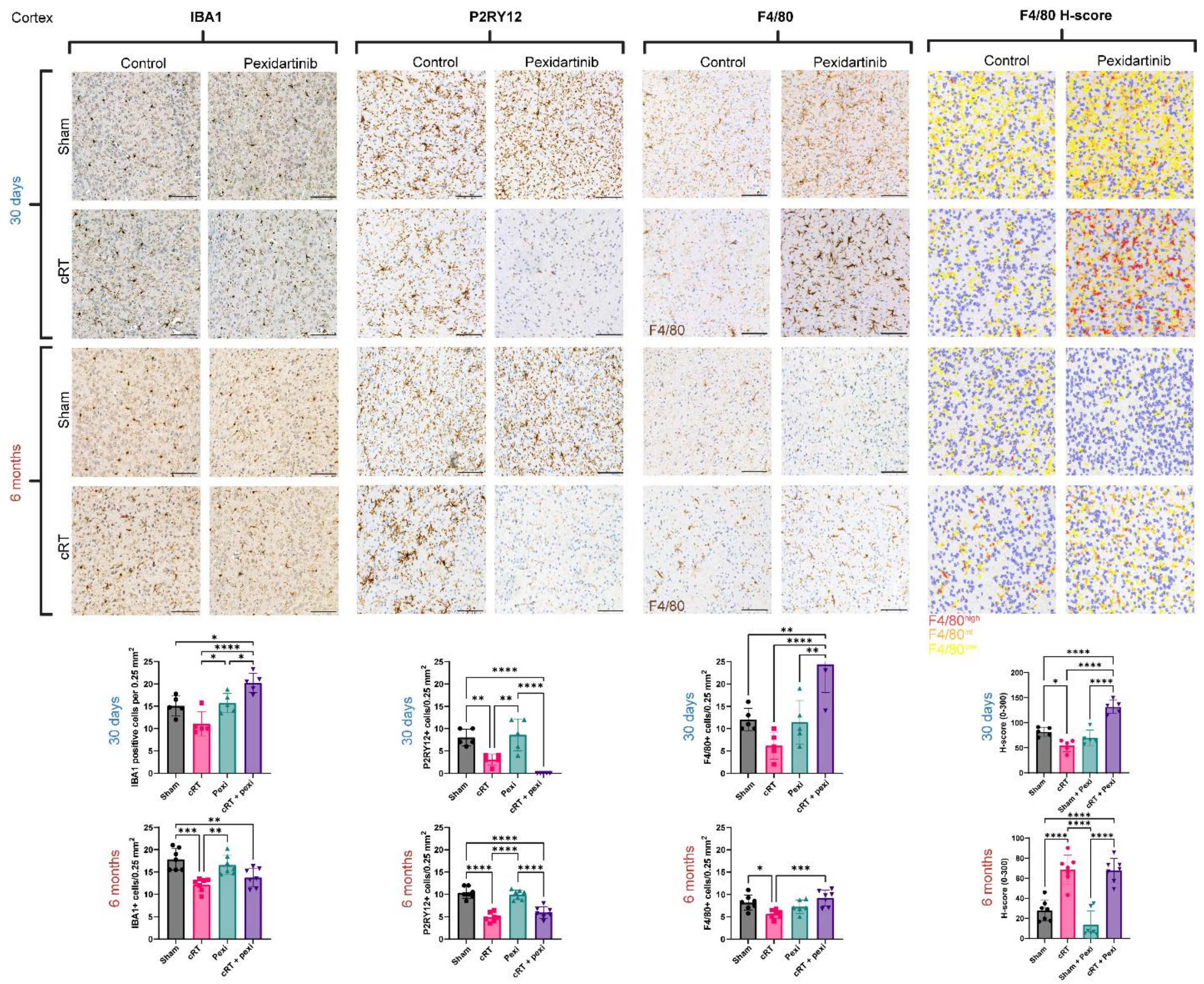


***Figure S2. IHC analysis of macrophage and microglial markers in the cortex.*** *Representative images of IHC and quantification on all three time points in the cortex. Each dot represents one animal, error bars represent standard deviation. * p<0.05, ** p<0.01, ***p<0.001, ****P<0.0001.*

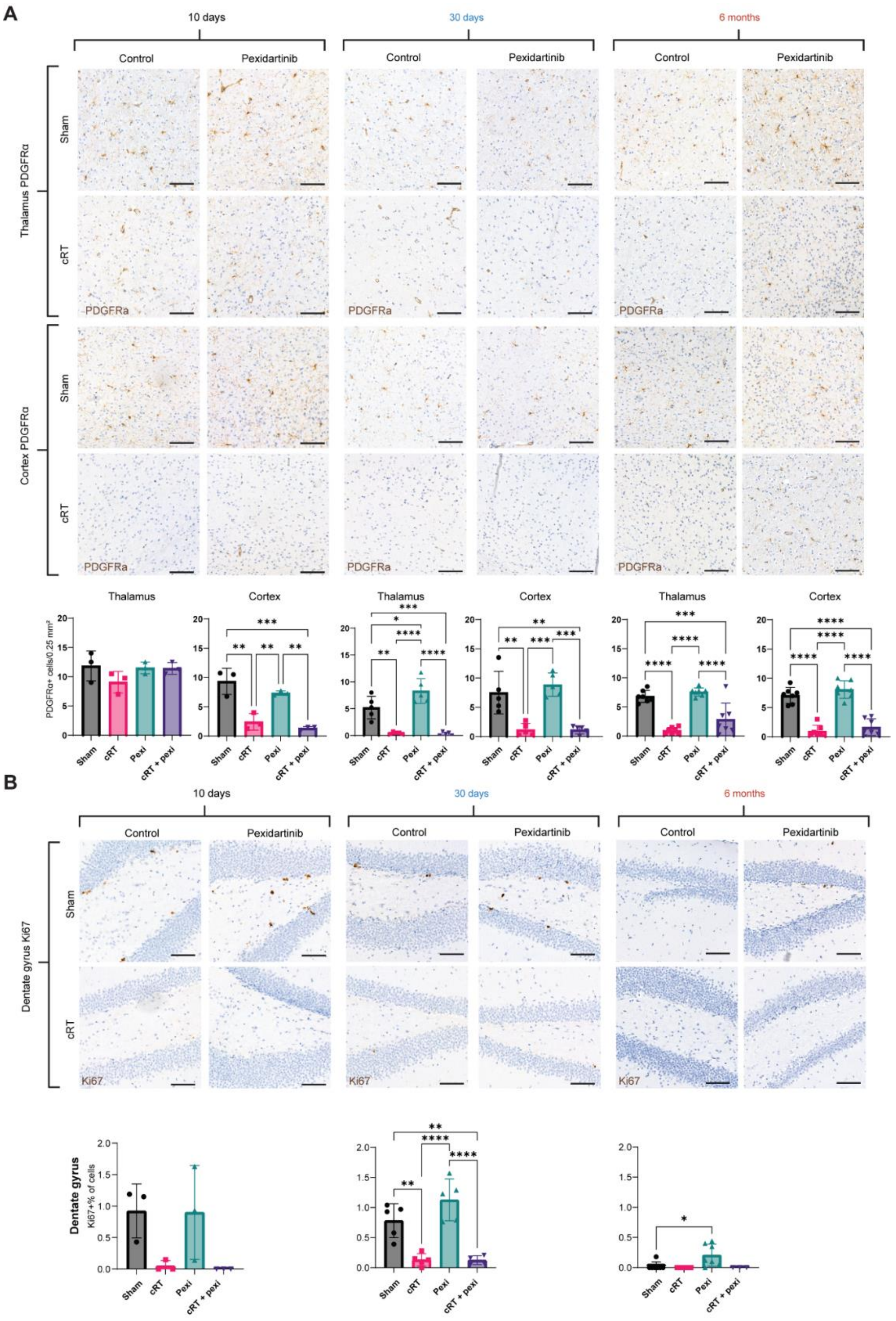


***Figure S3. IHC analysis of PDGFRa and Ki67.*** *A. Representative images of PDGFRa IHC and quantification on all three time points in the thalamus and cortex. B. Representative images of Ki67 IHC and quantification on all three time points in the dentate gyrus of the hippocampus. Each dot represents one animal, error bars represent standard deviation. * p<0.05, ** p<0.01, ***p<0.001, ****P<0.0001.*

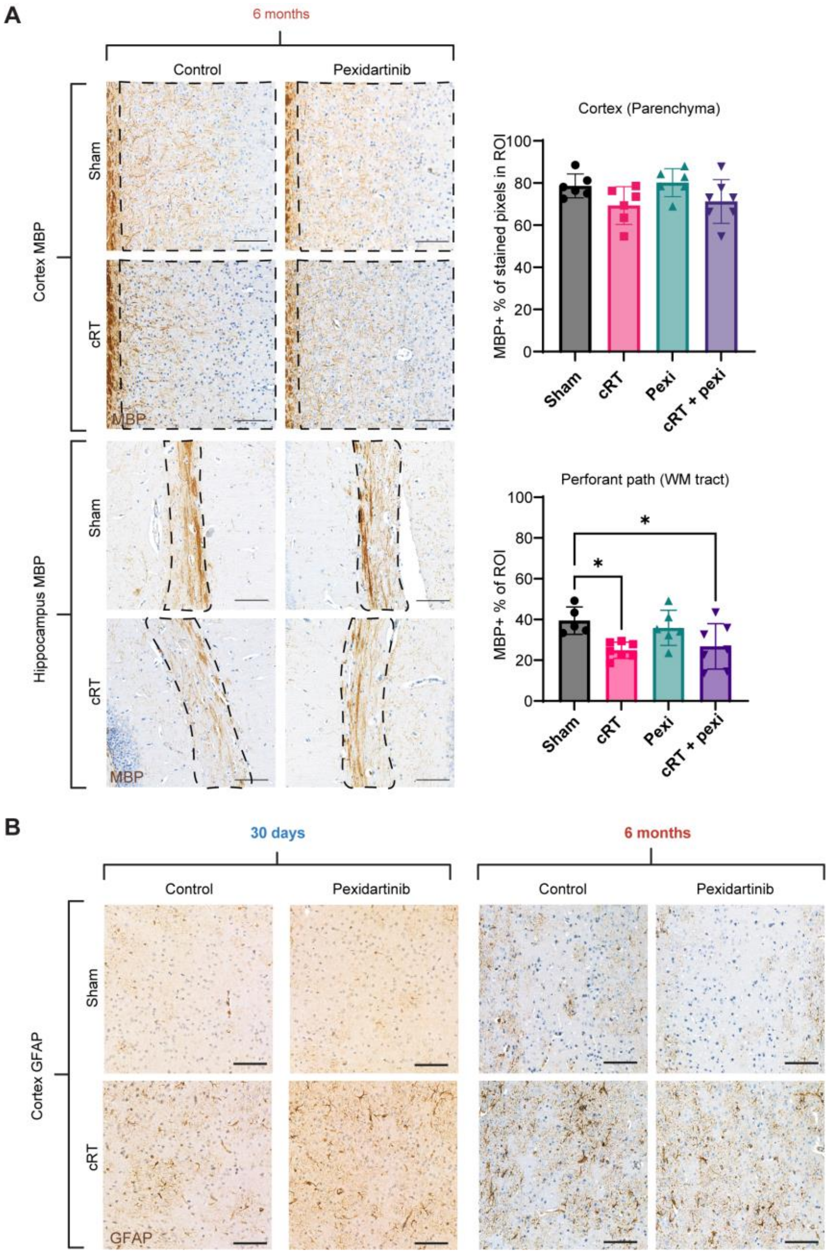


***Figure S4. IHC analysis of MBP and GFAP.*** *A. Representative images of MBP IHC and quantification in the cortex (parenchyma) and hippocampal perforant pathway (WM tract) at 6 months post cRT. Black dashed line represents ROI. Each dot represents one animal, error bars represent standard deviation. * p<0.05. B. Representative images of GFAP IHC at 30 days and 6 months post cRT in the cortex.*
